# Genome diversity and species richness in mammals

**DOI:** 10.1101/709311

**Authors:** John Herrick, Bianca Sclavi

**Author notes:** Corresponding author (B. Sclavi). Corresponding author at 3, rue des Jeûneurs, 75002 Paris, France. E-mail addresses (J. Herrick).

## Abstract

Evolutionary changes in karyotype have long been implicated in speciation events; however, the phylogenetic relationship between karyotype diversity and species richness in closely and distantly related mammalian lineages remains to be fully elucidated. Here we examine the association between genome diversity and species diversity across the class Mammalia. We tested five different metrics of genome diversity: clade-average genome size, standard deviation of genome size, diploid and fundamental numbers (karyotype diversity), sub-chromosomal rearrangements and percent synteny block conservation. We found a significant association between species richness (phylogenetic clade diversity) and genome diversity at both order and family level clades. Karyotype diversity provided the strongest support for a relationship between genome diversity and species diversity. Our results suggest that lineage specific variations in genome and karyotype stability can account for different levels of species diversity in mammals.

## Introduction

Genome size variation in eukaryotes spans over five orders of magnitude (Gregory, Nicol et al. 2007). In animals, genome size (C-value) ranges from 0.02 picograms (pg) in the nematode *Pratylenchcus coffeae* to over 120 picograms in lungfish and salamanders (Canapa, Barucca et al. 2015). Different animal taxa typically have characteristic ranges of genome size that can vary substantially between lineages and among groups. Salamanders, for example, have genome sizes ranging between 15 pg in the *Desmognathus* clade and 120 pg in the *Necturus* clade (Sessions 2008), whereas mammals have genome sizes ranging from 1.6 pg in bats to 9.2 pg in the red viscacha rat *(Tympanoctomys barrerae)* (Bromham 2011, Evans, Upham et al. 2017, Kapusta, Suh et al. 2017).

The large salamander genomes, and other giant genomes in plants and insects, have been attributed to a slow deletion rate of DNA (Bensasson, Petrov et al. 2001, Sun, López Arriaza et al. 2012, Kelly, Renny-Byfield et al. 2015), suggesting that these genomes are genetically more stable compared to taxa with smaller genomes. In the case of salamanders, the slow loss of DNA has been associated with significantly lower mutation rates in these organisms compared to other vertebrates (Mohlhenrich and Mueller 2016). Other studies have shown a similar negative relationship between genome size and mutation rates (Dores, Sollars et al. 1999, Herrick and Sclavi 2014), but the exact nature of the relationship, if any, remains to be elucidated.

Most of the variation in vertebrate genome size is attributable to differing amounts of non-coding DNA such as transposable elements, microsatellite DNA, LINES, SINES and pseudo genes (Feschotte and Pritham 2007, Graphodatsky, Trifonov et al. 2011, Metcalfe and Casane 2013, Bourque, Burns et al. 2018). These selectively neutral, or nearly neutral sequences are typically packaged into compact heterochromatin, which tends to be late replicating and devoid of actively expressed genes (Grewal and Moazed 2003, Grewal and Jia 2007, Aygün and Grewal 2010, Shermoen, McCleland et al. 2010). The function of heterochromatin in the genome remains controversial, while its contribution to adaptation and evolution is an open question (Nikolov and Taddei 2016, Allshire and Madhani 2018).

Nonetheless, species tend to differ in genomic terms more according to their noncoding DNA contents than according to their protein coding contents: once the amount of heterochromatic DNA is subtracted from total genomic DNA, all mammals have very similar and highly conserved genome sizes (Graphodatsky, Trifonov et al. 2011). Additionally, other studies have revealed a high degree of mammalian chromosome conservation (Ferguson-Smith and Trifonov 2007). Chromosome painting, which involves fluorescence *in situ* hybridization of a probe genome to a test genome (Balmus, Trifonov et al. 2007, Hua and Mikawa 2018), uncovered a high degree of homologous and syntenic regions among mammalian genomes (Ehrlich, Sankoff et al. 1997, Ferguson-Smith and Trifonov 2007, Graphodatsky, Trifonov et al. 2011, Zhao and Schranz 2019).

The genome of every species has a characteristic chromosome complement consisting of pairs of chromosomes that can be arranged according to size: the species karyotype (Sacerdot, Louis et al. 2018, Zhao and Schranz 2019). Although genome size diversity is relatively limited in mammals (spanning 4 to 5X), karyotype diversity is significantly less restricted (Graphodatsky, Trifonov et al. 2011, Kim, Farré et al. 2017). The smallest diploid mammalian karyotype is found in the Indian muntjac deer (2n = 6-7) (Yang, Carter et al. 1995). The largest, not surprisingly, is found in the red viscacha rat (2n = 102) (Evans, Upham et al. 2017). Hence, karyotype diversity is an order of magnitude larger than genome size diversity in mammals.

Karyotype diversity appears to be substantially greater in mammalian lineages with high levels of species diversity (Wilson, Sarich et al. 1974, Bush, Case et al. 1977, Bengtsson 1980, Maruyama and Imai 1981). This pattern is evident both across different taxa such as salamanders, frogs, mammals and reptiles and within different lineages (Bush, Case et al. 1977, Bengtsson 1980, Olmo 2005, Olmo 2006). The mouse genome, for example, is much more rearranged than that of most other taxa in the class Mammalia, and the order Rodentia comprises more than 40% of all mammalian species (Graphodatsky, Trifonov et al. 2011, Romanenko, Perelman et al. 2012). Moreover, one third of Rodentia species belong to a single superfamily, the Muroidea, while other lineages of Rodentia are relatively species poor. These observations indicate unequal and widely varying rates of genome and species evolution in the different mammalian lineages despite the highly conserved coding complement of DNA (Bromham, Hua et al. 2015, Kapusta, Suh et al. 2017).

The ability to detect and measure karyotype differences has increased enormously over the past twenty years (Payseur and Rieseberg 2016, Hua and Mikawa 2018), resulting in a rapidly growing collection of high resolution datasets that allow detailed analyses of the relationships between genome evolution and other evolutionary variables including physiological traits (body size, metabolic rate), life-history traits (r vs K-strategists) and ecological traits (geographic ranges, niche rate, diversification rate and species diversity) (Castro-Insua, Gómez-Rodríguez et al. 2018).

Recently, Martinez and co-workers investigated karyotype diversification rates in mammals and tested three different models of speciation: the metabolic rate hypothesis, the reproductive rate hypothesis, and the geographic range hypothesis (Martinez, Jacobina et al. 2017). They concluded that the geographic range hypothesis, according to which larger geographic ranges favor fixation of DNA rearrangements, best explains rates of karyotype diversity in mammals: high diversity is strongly associated with correspondingly large geographic area. The phylogenetic relationship between species richness and genome diversity (genome size and karyotype diversities), however, was not specifically addressed in their study.

In the following, we report our findings on the phylogenetic relationship between species richness (phylogenetic clade diversity) and genome diversity. We employed five different metrics to assess the co-variation between species richness and genome diversity: average genome size in a lineage, standard deviation of genome size, percent conserved of synteny blocks, and frequencies of sub-chromosomal rearrangements and gross genomic differences in diploid number and number of chromosomal arms (fundamental number). Our findings indicate a significant and strong relationship across mammalian lineages between genome stability and respective species diversity in each of the corresponding lineages.

## Materials and methods

### C-values

Genome sizes were obtained from the Animal Genome Size Database (Gregory 2015). C-value refers to haploid nuclear DNA content (1C). Reported polyploids, when indicated in the Animal Genome Size Database, were removed from the analyses. Average C-values were determined for each species when more than one C-value is recorded in the database. These values were used to calculate the average C-value of each family-level clade and each order-level clade.

### Karyotype diversity

The data on family-level clade time of divergence, rKDmacro, rKDmicro (respectively macro and micro-structural rate of karyotype diversification), and karyotype diversity (KD) were obtained from Martinez *et al.* 2017. They determined rKDmacro from the number of different diploid number of chromosomes (2n) and the fundamental number (Fn) combinations in each family level clade and normalized this number by the family time of origin to obtain the rate. Here, the same data was used to determine rKDmacro at the order level. The radiation time at the order level was obtained from TimeTree (www.timetree.org). From different bibliographic sources, Martinez *et al.* determined rKDmircro from chromosome painting and chromosome banding of 208 mammalian species. The rKDmacro and rKDmicro rates quantify the number of chromosome changes per million years since the most recent common ancestor of the family clade.

### Phylogenetic tree

The phylogenetic ML tree of mammals was created by Meredith *et al.* on the basis of the amino acid matrix of 164 species rooted with five vertebrate outgroups (zerbrafish, green anole, chicken, zebrafish and frog). The Meredith *et al* family level tree was used to obtain an order level tree (Meredith, Janečka et al. 2011). Species were assigned to families and families to orders at first using the *taxize* package in R (Chamberlain and Szöcs 2013). These were then manually verified against the taxonomy of the Meredith tree. The HighLevelTree function in the *EvobiR* package in R by Heath Blackmon was used to obtain the order level tree (evobiR: evolutionary biology in R. R package version 1.0. http://CRAN.R-project.org/package=evobiR).

### Species diversity

Species diversity for family-level clades was obtained from Castro-Insua *et al.* 2018.

### Synteny analysis

The average percentage gene synteny conservation at the family and order level was obtained from the data of Zhao *et al.* 2019. They determined the percentage synteny conservation for 87 species yielding 7569 whole genome comparisons of protein-coding regions for microsynteny block detection. They used the number of syntenic pairs plus the number of colinear tandem genes relative to the number of all annotated genes to determine each pairwise syntenic percentage. Here, only data with N50 values above 2 were used for the analyses.

### Regression analysis

The *pgls* analysis was carried out in R with the *caper* package. The maximum likelihood value of lambda was allowed to vary while kappa and delta were set to 1 as in Kozak and Wiens (Kozak and Wiens 2016). For the *pgls* analysis, we used the ML family level tree from Meredith *et al.* and the order level tree described above.

## Results

### Uneven distribution of species richness is evident in the class Mammalia

Examination of the phylogenetic tree for the class Mammalia at the order and family levels reveals a remarkable degree of heterogeneity in species richness ranging from just five living species in the order Monotreme to up to 2500 living species in the order Rodentia (Figure 1A and B) (Purvis and Hector 2000, Jones and Safi 2011, Safi, Cianciaruso et al. 2011, Davis, Faurby et al. 2018). This observation is evident at the lower phylogenetic level of genus (not shown), indicating that species richness varies substantially between the different lineages throughout the Mammalia phylogenetic tree (Castro-Insua, Gómez-Rodríguez et al. 2018, Upham, Esselstyn et al. 2019).

**Figure 1A:**
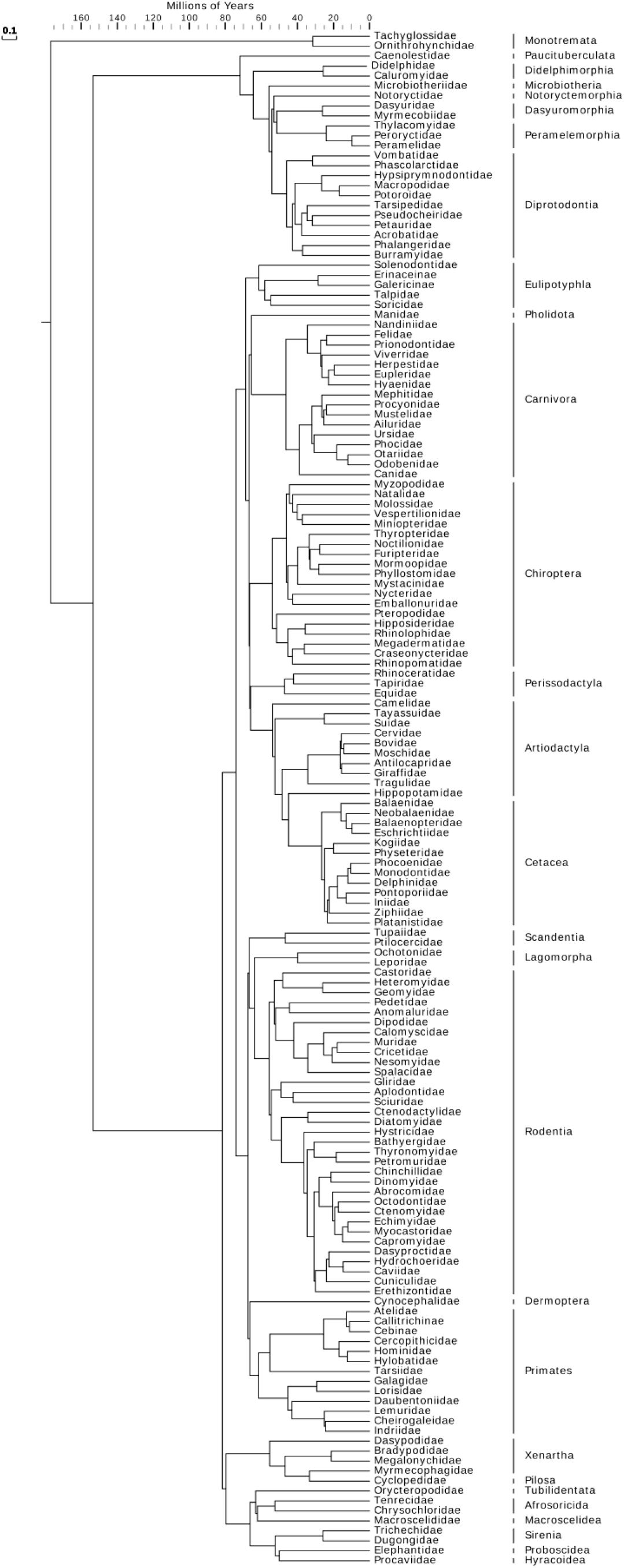
Phylogenetic tree of the family level clades from Meredith *et al.* 2011.

**Figure 1B.**
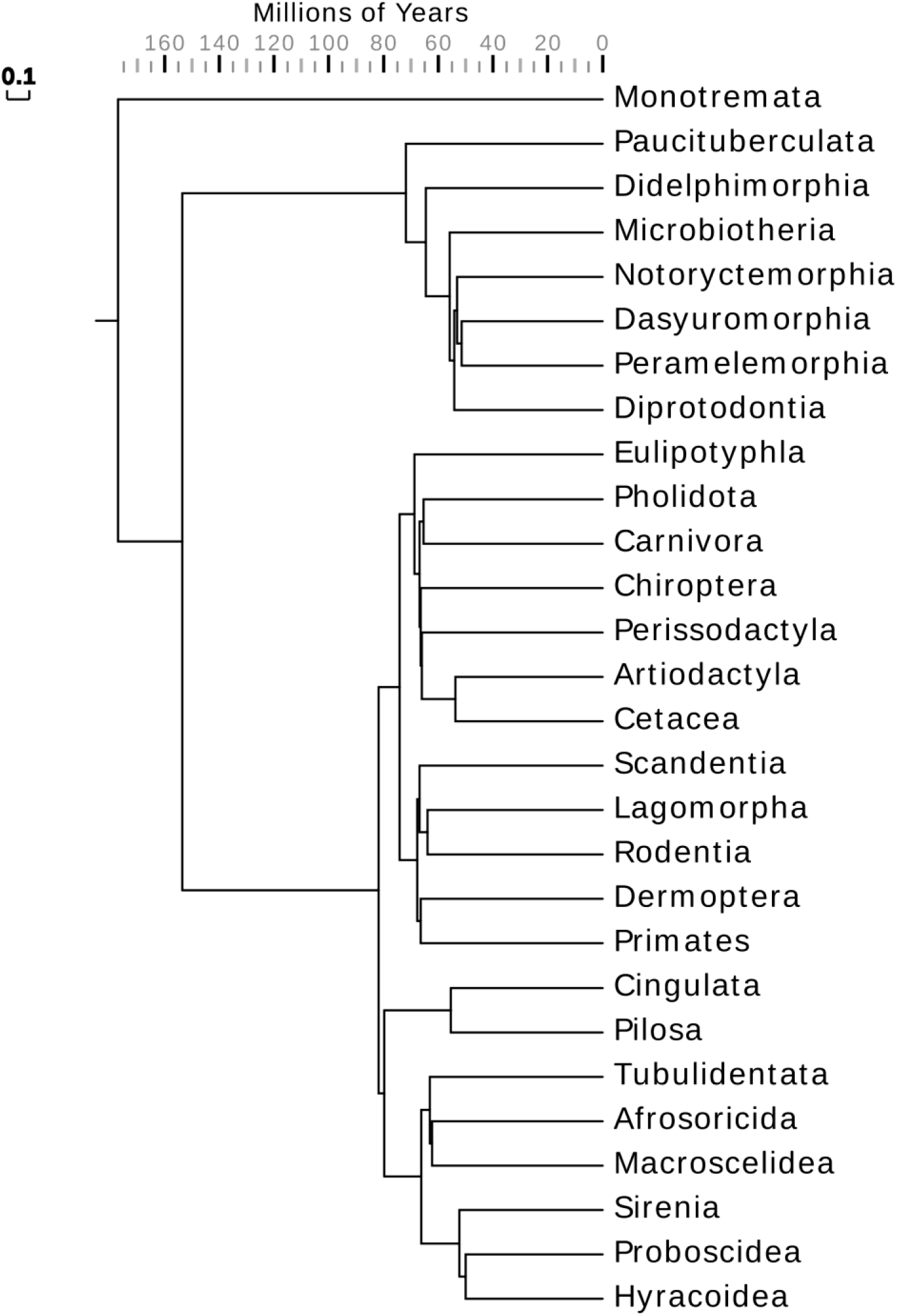
Phylogenetic tree of the order level clades derived from Meredith *et al.* 2011.

Muroids present a striking example of this unevenness. They comprise an estimated 1,620 species split between three lineages representing some 30 *%* of mammalian species richness (about 5,400 defined species). The depauperate Platacanthomyidae containing two monotypic genera split from the nineteen other mostly species-rich murid subfamilies about 45.2 million years ago (Mya). (Isaac, Jones et al. 2005, Steppan and Schenk 2017).

### Uneven distributions of karyotype diversity at order and family level clades

Figure 2A shows box plots of genome size in mammals for order level clades (Gregory 2015). The coefficient of variation of C-value (CV of C-value) provides an estimate of genome size diversity relative to average genome size in a clade (Hardie and Hebert 2004, Sclavi and Herrick 2019). A trend between CV of C-value and average C-value is not apparent in the mammalian phylogenetic tree (not shown), as expected from the relatively narrow range of genome sizes. At the order level, the smallest CV of C-value was 0.02 in Dasyuromorphia (mean C-value = 3.43 pg) and the largest was 0.21 in the Eulipotyphyla (mean C-value = 3.06). At the family level, the smallest CV of C-value is 0.005 in the Callitricidae (mean C-value = 4.02 pg) and the largest is 0.21 in the Geomyidae (mean C-value = 3.62 pg).

**Figure 2A.**
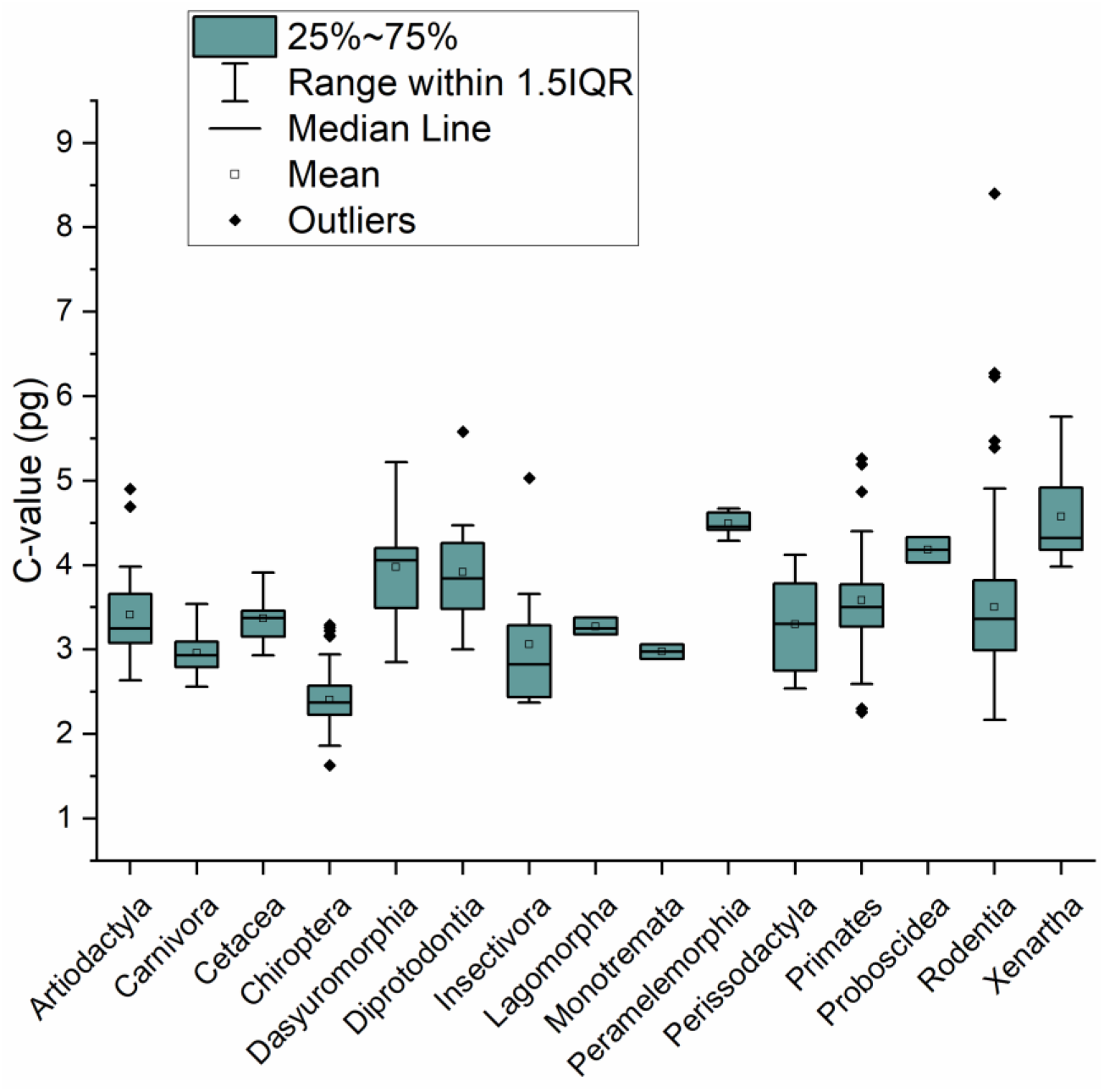
Distribution of genome size within each of the order level clades. Genome sizes in pg were obtained from the Animal Genome Size Database (Gregory, T. R. 2015).

Karyotype diversity (KD; see below), in contrast to C-value and CV of C-value, exhibits a level of unevenness similar to the uneven distribution found for species richness (SR) across the Mammalia orders (Figure 2B). With the exception of Chiroptera and Peramelemorphia, species richness increases consistently with karyotype diversity: Rodentia, the most species diverse order, has the highest levels of karyotype and species diversities, while Tubilidentata has the lowest. The extreme unevenness and similar shapes of the distributions of karyotype diversity and species diversity suggest that KD and SR are diverging similarly and in pace from the last common ancestor in each clade. On average, 13 speciation events occur per change in karyotype (Figure 2C; slope = 0.69).

**Figure 2B.**
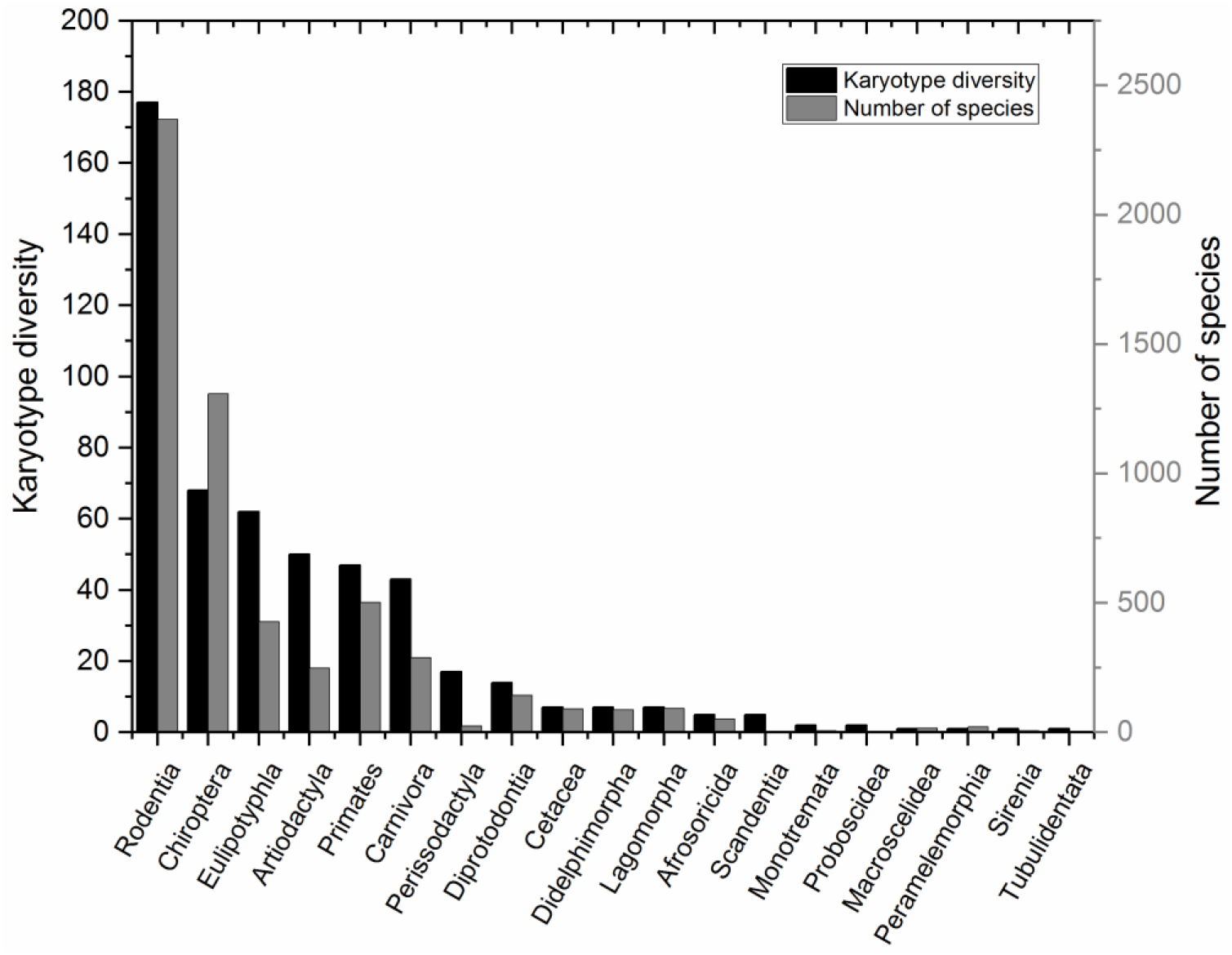
Karyotype diversity (left y-axis) and species richness (right y-axis) for the order level clades.

**Figure 2C.**
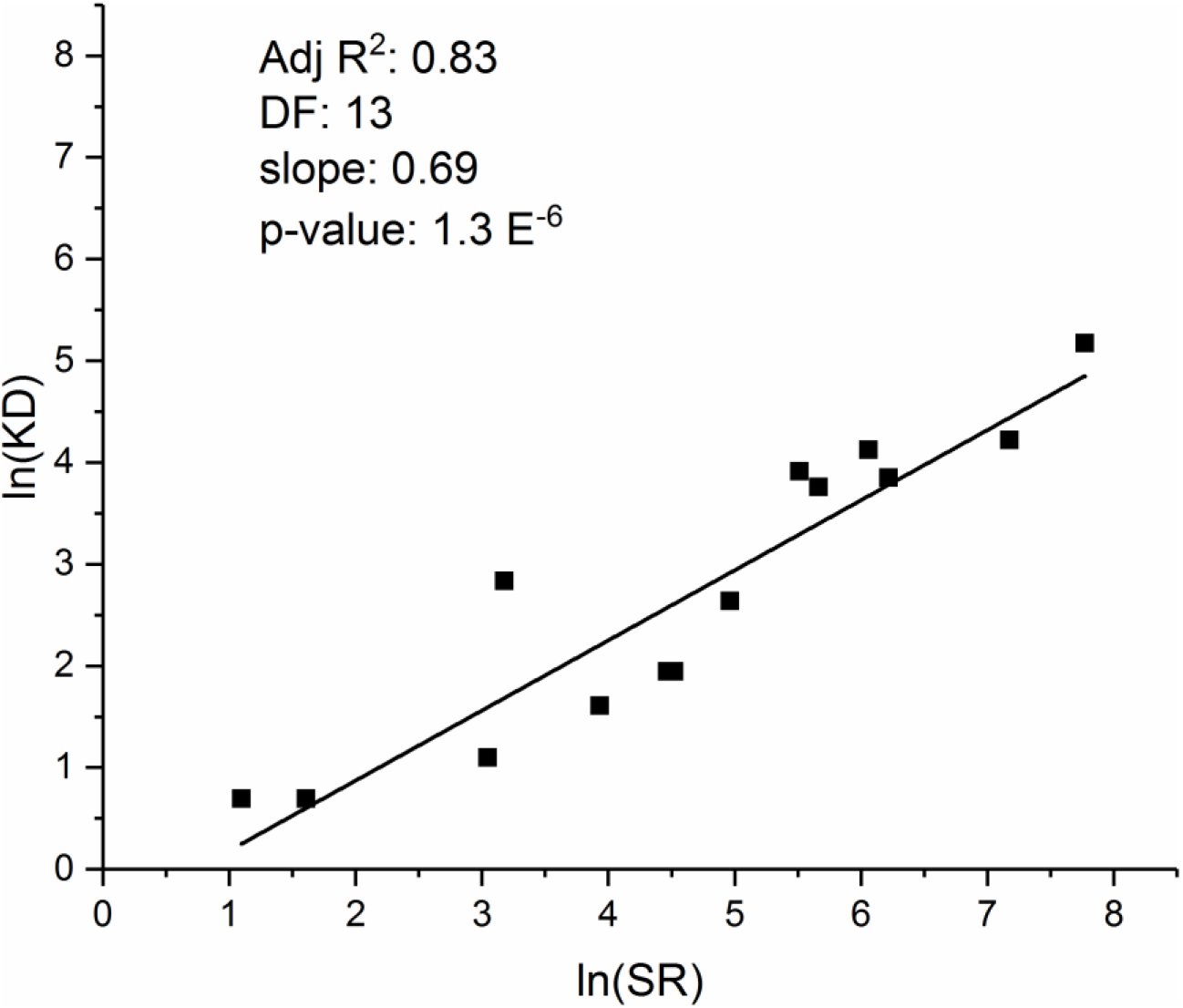
Karyotype diversity vs. species richness for the order level clades.

### Stem age versus average C-value: a junk DNA molecular clock?

We next examined the evolution in genome size in time by regressing clade stem age to clade average genome size at the family level clades using ordinary least squares (OLS). Figure 3 reveals two distinct populations: before and after the Cretaceous-Tertiary (KT) boundary. A constant linear increase with time in clade average genome size begins at the KT boundary 60 million years ago (Mya), confirming a clear linear dependence on time (R^2^ = 0.30; p = 4 x 10^-6^). Changes in mean C-value with time suggest a rate of increase on the order of 0.028 pg/Mya (y-intercept = 4.2 pg).

**Figure 3.**
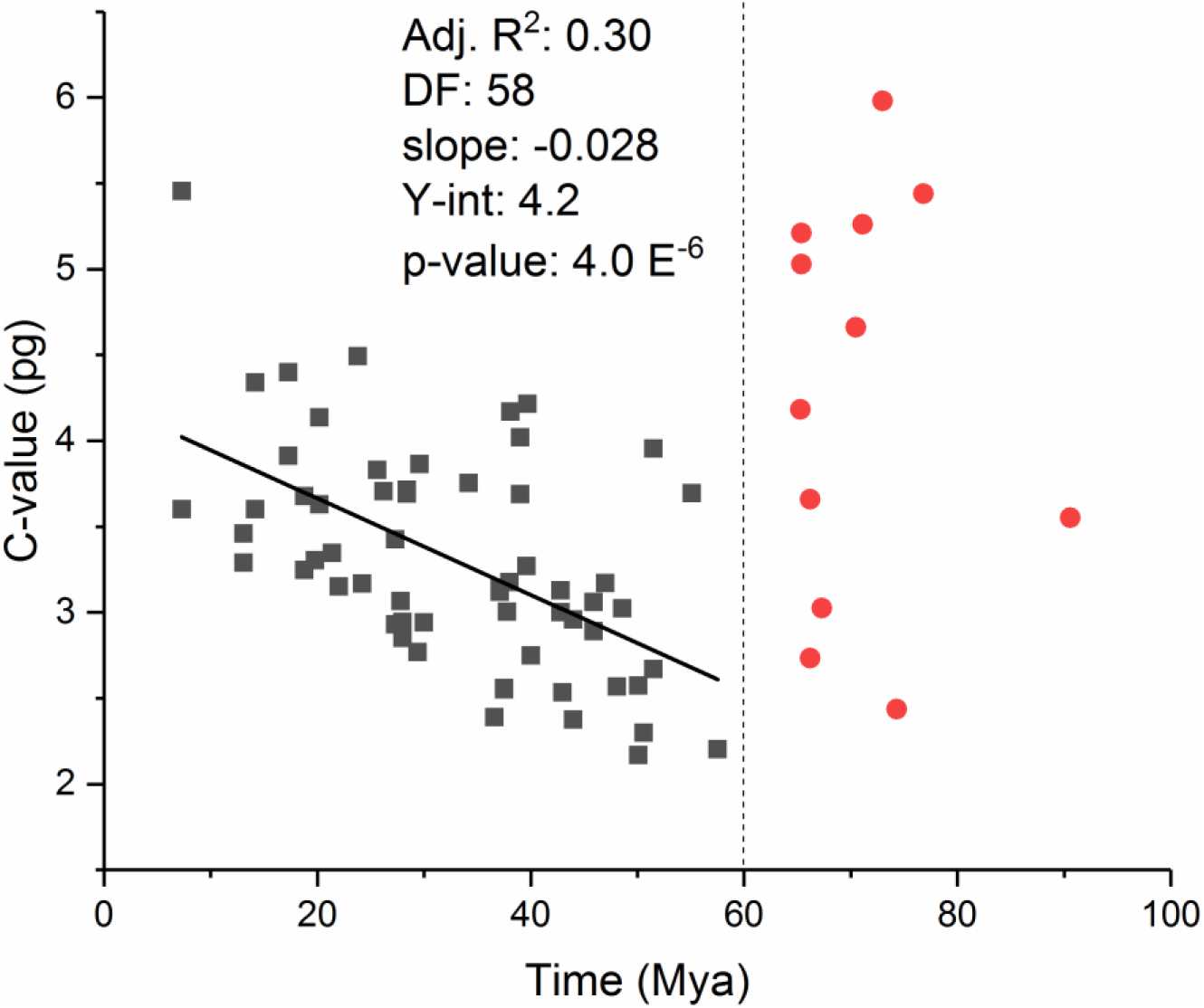
Average family level clades C-value as a function of the time of origin. The dotted line indicates the K-T boundary.

In striking contrast, extant lineages originating before 60 Mya showed no correlation between time and C-value (R^2^ = 0.002). The median C-value in the group belonging to the pre-KT boundary group is about 4.42 pg (range = 2.4 to 5.98 pg); the median in the post-KT group, for the 40 to 60 Mya time window, is about 2.82 pg (range = 2.17 to 3.95 pg), consistent with a decrease in C-value at the KT boundary (Rho, Zhou et al. 2009, Grossnickle, Smith et al. 2019). The observed change in genome size through time beginning 60 Mya is suggestive of a molecular clock in mammals, at least since the KT extinction event (Bulmer, Wolfe et al. 1991, Trusov and Dear 1996), though relative rates differ substantially between lineages (Goldie, Lanfear et al. 2011, Bromham, Hua et al. 2015, Moorjani, Amorim et al. 2016).

### C-value diversity and species richness at the order and family levels

The pattern of average C-value variation and the large heterogeneity in species richness among mammalian clades observed in Figures 1 and 2 suggests a potential relationship between C-value and species richness (SR). Phylogenetic least squares analyses (pgls), however, revealed a weak and only marginally significant correlation at the order level clades (R^2^ = 0.14; p = 0.06), and no correlation between C-value and SR at the family level (R^2^ = 0.004; p = 0.28; Table 1 and 2), consistent with average C-values being substantially more constrained compared to species richness.

**Table 1.** Results of pgls analysis of the order level clades.

|  | *Adj R<sup>2</sup>* | *p* | *AICc* | *Pagel's λ* | *F* | *DF* | *Slope* |
| --- | --- | --- | --- | --- | --- | --- | --- |
| ***Ln(SR) vs log(KD)*** | **0.84** | **2.1 x 10<sup>-6</sup>** | **35** | **0** | **71** | **12** | **+** |
| ***Ln(SR) vs log(rKDmacro)*** | **0.83** | **2.9 x 10<sup>-6</sup>** | **36** | **0** | **67** | **12** | **+** |
| ***Ln(SR) vs log(rKDmicro)*** | **0.45** | **0.03** | **36** | **0** | **7.5** | **7** | **+** |
| *Ln(SR) vs synteny* | 0.07 | 0.19 | 56 | 1 | 1.9 | 11 | - |
| ***Ln(SR) vs CV of C-value*** | **0.38** | **0.008** | **58** | **0** | **9.8** | **13** | **+** |
| ***Ln(SR) vs SD of C-value*** | **0.31** | **0.02** | **59** | **0** | **7.3** | **13** | **+** |
| *Ln(SR) vs log(C-value)* | 0.13 | 0.07 | 76 | 1 | 3.6 | 17 | - |
| ***SR vs KD*** | **0.90** | **1.7 x 10<sup>-7</sup>** | **193** | **0** | **114** | **12** | **+** |
| ***Log(rKDmacro) vs log(rKDmicro)*** | **0.43** | **0.02** | **16** | **0** | **7.8** | **8** | **+** |
| *Log(rKDmacro) vs synteny* | 0.1 | 0.18 | 21 | 0 | 2 | 9 | - |
| *Log(rKDmicro) vs synteny* | -0.13 | 0.8 | 1.1 | 0 | 0.07 | 7 | - |

**Table 2.**
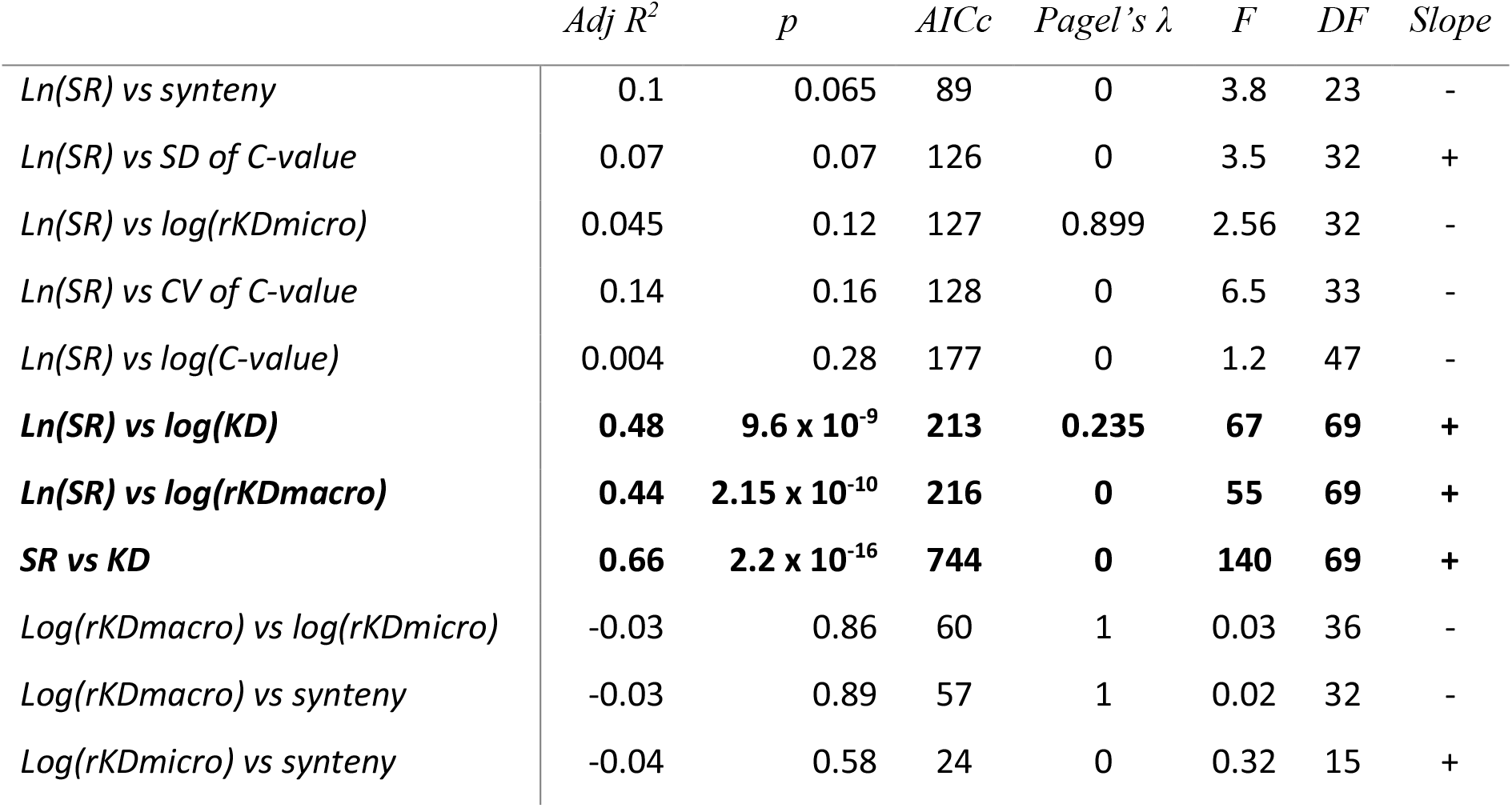
Results of pgls analysis of the family level clades.

|  | *Adj R<sup>2</sup>* | *p* | *AICc* | *Pagel's λ* | *F* | *DF* | *Slope* |
| --- | --- | --- | --- | --- | --- | --- | --- |
| *Ln(SR) vs synteny* | 0.1 | 0.065 | 89 | 0 | 3.8 | 23 | - |
| *Ln(SR) vs SD of C-value* | 0.07 | 0.07 | 126 | 0 | 3.5 | 32 | + |
| *Ln(SR) vs log(rKDmicro)* | 0.045 | 0.12 | 127 | 0.899 | 2.56 | 32 | - |
| *Ln(SR) vs CV of C-value* | 0.14 | 0.16 | 128 | 0 | 6.5 | 33 | - |
| *Ln(SR) vs log(C-value)* | 0.004 | 0.28 | 177 | 0 | 1.2 | 47 | - |
| ***Ln(SR) vs log(KD)*** | **0.48** | **9.6 x 10<sup>-9</sup>** | **213** | **0.235** | **67** | **69** | **+** |
| ***Ln(SR) vs log(rKDmacro)*** | **0.44** | **2.15 x 10<sup>-10</sup>** | **216** | **0** | **55** | **69** | **+** |
| ***SR vs KD*** | **0.66** | **2.2 x 10<sup>-16</sup>** | **744** | **0** | **140** | **69** | **+** |
| *Log(rKDmacro) vs log(rKDmicro)* | -0.03 | 0.86 | 60 | 1 | 0.03 | 36 | - |
| *Log(rKDmacro) vs synteny* | -0.03 | 0.89 | 57 | 1 | 0.02 | 32 | - |
| *Log(rKDmicro) vs synteny* | -0.04 | 0.58 | 24 | 0 | 0.32 | 15 | + |

In contrast to C-value, pgls analysis revealed a relatively strong and significant correlation between CV of C-value and SR at the order level clades in Mammalia (R^2^ = 0.38, p = 0.008; Table 1). A much weaker and insignificant correlation was found at the family level (R^2^ = 0.14; p = 0.16; Table 2). Because the mean genome size of mammals is restricted in range, we examined the absolute C-value diversity as measured by standard deviation. pgls analysis established a significant correlation between SR and standard deviation of C-value at the order level clades but not at the family level (R^2^ = 0.31; p = 0.02; R^2^ = 0.07, p = 0.07; Tables 1 and 2).

### Karyotype diversity and species richness at the order and family levels

Martinez *et al.* developed two quantitative metrics of karyotype diversity (Martinez, Jacobina *et al.* 2017): rKDmacro and rKDmicro. rKDmacro measures gross karyotype diversity in terms of diploid chromosome number (2n) and the number of chromosomal arms, or fundamental number (Fn). rKDmicro measures the number of inter and intra-chromosomal rearrangements obtained from chromosomal painting. Additionally, Zhao *et al.* have recently reported a study on mammalian synteny blocks. In this study, they measured the percent synteny conserved among 87 mammalian species (Zhao and Schranz 2019).

pgls analysis of rKDmacro versus SR revealed very strong and highly significant correlations at both order and family level clades (R^2^ = 0.83, p = 2 x 10^-6^; R^2^ = 0.44, p = 2.15 x 10^-10^; Tables 1 and 2). We then converted rKDmacro, a phylogenetic rate metric, into a time-independent karyotype diversity measure (KD) to assess the total karyotype diversity and its relationship to SR. pgls analysis of the non-logged KD metric versus SR provided the strongest correlations at both order and family levels (R^2^ = 0.90, p = 1.7 x 10^-7^; R^2^ = 0.66, p = 2.2 x 10^-16^; Table 1 and 2). Consistent with rKDmacro correlating with SR, our pgls examination of rKDmicro versus SR showed strong, significant correlations at the order level clades, but much weaker correlations at the family level clades (R^2^ = 0.45, p = 0.03 and R^2^ =0.045, p = 0.12 respectively; Tables 1 and 2). This observation is consistent with rKDmacro and rKDmicro being correlated at the order (R^2^ = 0.43; p = 0.02; Table 1) but not at the family levels (R2 = −0.03, p = 0.86; Table 2).

Our pgls analysis of conserved synteny blocks and SR revealed either no correlation at the order level or only a weak and marginally significant correlation at the family levels (R^2^ = 0.07, p = 0.19 and R^2^ = 0.1, p = 0.065 respectively; Tables 1 and 2). Likewise, synteny is not correlated with rKDmacro either at the order level or at the family level (Tables 1 and 2). No significant correlations were found between synteny conservation and other metrics of genome diversity (C-value, rKDmicro, SD of C-value and CV of C-value). Together, these findings support previous reports of synteny being highly conserved across the class Mammalia (Ferguson-Smith and Trifonov 2007, Graphodatsky, Trifonov et al. 2011).

## Discussion

We previously reported significant relationships between species richness and both C-value and genome size diversity (CV of C-value) in salamanders (Sclavi and Herrick 2019). Here we confirm and extend those findings to genome size diversity and species richness in mammals. Since salamanders have significantly larger variation in taxonomic average C-values compared to mammals, we employed two additional metrics of genome diversity developed by Martinez *et al:* rKDmacro and rKDmicro, which measure phylogenetic rates of karyotype diversity at the genome and sub-chromosomal levels. We also examined absolute karyotype structural diversity using the KD metric, which provided the strongest and most significant evidence that differences in karyotype diversity, and hence genome diversity, contribute significantly to explain differences in species diversity in Mammalia.

Our finding on the unevenness of KD distributions provides an additional explanation to the extreme unevenness of the SR distribution in the mammalian phylogenetic tree (Figure 2B)(Davis, Faurby et al. 2018, Grossnickle, Smith et al. 2019, Upham, Esselstyn et al. 2019). Our findings also confirm a dramatic change in genome size distributions in mammals occurring at the KT boundary (Rho, Zhou et al. 2009, Grossnickle, Smith et al. 2019). Genome sizes appear to have decreased significantly at the KT boundary, and then steadily increased with ensuing speciation events over the following 60 My: older extant species that emerged at the KT boundary have approximately 2X less DNA than younger extant species that emerged more recently (Rho, Zhou *et al.* 2009).

The two metrics, rKDmacro and rKDmicro, measure cytogenetically separable modifications of karyotypes. rKDmacro measures changes in chromosome and fundamental numbers at the genomic level, whereas rKDmicro measures sub-chromosomal rearrangements (SCR) that do not significantly affect the overall cytogenetic organization of the genome. We suggest that these different cytogenetic levels of structural change reflect two different modes of genome dynamics and instability: rKDmacro reflects double strand breaks in DNA (DSB) and Robertsonian translocations (Rb) that likely occur during meiosis (Pardo-Manuel de Villena and Sapienza 2001, Graphodatsky, Trifonov et al. 2011, Romanenko, Perelman et al. 2012), while rKDmicro reflects DSBs that occur for the most part during interphase of the germline cell cycle (Mao, Bozzella et al. 2008, Mao, Bozzella et al. 2008, Shrivastav, De Haro et al. 2008, Shibata, Conrad et al. 2011, Ambrosio, Di Palo et al. 2016, Muramoto, Oda et al. 2018). This hypothesis, however, remains to be further examined.

The observation that these two modes of instability differentially impact species diversity is highly suggestive: Rb and chromosomal loss/gain evidently result in stronger reproductive isolation and possibly stronger barriers to introgression compared to SCRs, and therefore they might play a more significant role in the mechanisms initiating and consolidating speciation events (Dion-Côté and Barbash 2017). The different impact of each mode on species diversity might also account for the highly conserved nature of the mammalian genome (low levels of SCR and high synteny block conservation) despite the highly variable karyotype diversity (high levels of Rb and chromosome loss/gain). Together, our observations support the chromosome speciation hypothesis; and establish a molecular link between chromosome speciation and the geographic range hypothesis, which is an ecological model of speciation proposing that larger geographical ranges promote the fixation of rearrangements generated by genome instability (Faria and Navarro 2010, Potter, Bragg et al. 2017).

Interestingly, karyotype diversity does not seem to be relevant in terms of phenotype as exhibited by the large difference in karyotype but close phenotypic resemblance between Chinese (2n = 46) and Indian munctjacs (2n = 6-7) (Ferguson-Smith and Trifonov 2007). Likewise, Rb translocations have led to the identification of over 40 races of the house mouse *Mus musculus domesticus* worldwide, which are otherwise phenotypically very similar (Britton-Davidian, Catalan et al. 2000, Britton-Davidian, Catalan et al. 2005, Garagna, Page et al. 2014). These observations suggest that morphological diversification is uncoupled from speciation events associated with changes in karyotype diversity (Venditti, Meade et al. 2011).

Our findings suggest that karyotype changes (rKDmacro and rKDmicro) might precede other mutational events that later result in speciation (gene specific mutations), or conversely these different mutational modes might occur independently of each other but in parallel (Venditti and Pagel 2010). Our finding that thirteen speciation events occur on average per change in karyotype supports the proposal that these evolutionary events (karyotype diversification, mutation and speciation) can proceed in parallel in a manner that is mutually reinforcing during the speciation process (Figure 2C). The absence of a correlation between of synteny (gene order) and either SR or KD, however, suggests that SR and speciation depend more on genome stability at the macro level than at the gene or gene order level (Tables 1 and 2).

The differential relationship between the two different forms of karyotype modification (rKDmacro vs. rKDmicro) and species richness addressed here supports the hypothesis that non-coding DNA is of structural importance cytogenetically in driving the process of speciation, with differing amounts of heterochromatin-associated non-coding DNA accounting, in part, for differing amounts of species richness. Karyotype diversity (KD and rKDmacro), which largely accounts for species-dependent differences in C-value, exhibits the strongest correlations with species richness. Clearly, the relationship between heterochromatin content and species richness merits further attention.

We reported in salamanders the results of a path analysis that showed genome diversity and species richness appear to evolve independently and in parallel over time (Sclavi and Herrick 2019). Our findings in salamanders and mammals therefore raise an outstanding question: Does higher genome diversity merely reflect the higher species diversity in the different taxa (more species per taxon entailing more karyotype diversity per taxon)? In other words, are our observations artefacts due to a sampling bias?

This seems unlikely, since karyotype diversity should, in that case, strongly associate with phylogenetic diversity (species richness) at all phylogenetic levels. This is clearly not the case. The very strong correlations at the order level weaken substantially at the family level (Tables 1 and 2), suggesting that the relationship between genome diversity and species richness depends more on phylogenetic distance (time) than standing species richness in a clade.

Moreover, the association between karyotype diversity and SR would be expected to be linear and directly proportional if KD merely reflected SR in all taxa at all clade levels. This too is not the case: up to thirteen speciation events occur per change in karyotype, indicating that sources other than SR contribute to the relationship between these two phylogenetic variables. The relationship between genome diversity and species richness is therefore more likely attributable to an unknown underlying evolutionary process rather than to statistical happenstance or taxonomic sampling bias.

Alternatively, our observations elicit related but different questions: is there a mechanistic relationship operating at the *molecular level* of genome stability such that varying rates of genome diversification differentially promote propensities to speciate in the different taxonomic lineages (Wilson, Sarich et al. 1974, Bush, Case et al. 1977, Bengtsson 1980, Herrick 2011, Herrick 2011, Sclavi and Herrick 2019)? Can such a mechanism, underlying genome stability, account for the extreme species unevenness apparent in many of the vertebrate lineages? If so, what then is that mechanism? We will address those and other questions in an accompanying paper that examines the mechanisms governing genome stability and how those mechanisms might influence speciation and species diversity in mammals and other eukaryotes.

